# Conditional Freezing, Flight and Darting?

**DOI:** 10.1101/2021.12.02.470975

**Authors:** Jeremy M. Trott, Ann N. Hoffman, Irina Zhuravka, Michael S. Fanselow

## Abstract

Fear conditioning is one of the most frequently used laboratory procedures for modeling learning and memory generally, and anxiety disorders in particular. The conditional response (CR) used in the majority of fear conditioning studies in rodents is freezing. Recently, it has been reported that under certain conditions, running, jumping or darting replaces freezing as the dominant CR. These findings raise both a critical methodological problem and an important theoretical issue. If only freezing is measured but rodents express their learning with a different response, then significant instances of learning, memory, or fear may be missed. In terms of theory, whatever conditions lead to these different behaviors may be a key to how animals transition between different defensive responses and different emotional states. We replicated these past results but along with several novel control conditions. Contrary to the prior conclusions, running and darting were entirely a result of nonassociative processes and were actually suppressed by associative learning. Darting and flight were taken to be analogous to nonassociative startle or alpha responses that are potentiated by fear. On the other hand, freezing was the purest reflection of associative learning. We also uncovered a rule that describes when these movements replace freezing: When afraid, freeze until there is a sudden novel change in stimulation, then burst into vigorous flight attempts. This rule may also govern the change from fear to panic.

## Introduction

Fear limits the behaviors available to an animal to its species-specific defense reactions (SSDRs), thereby precluding more flexible voluntary behavior (Bolles, 1970). This characteristic is one reason that conditions characterized by high fear levels such as anxiety disorders are so maladaptive (Fanselow, 2018). It is also one reason that Pavlovian fear conditioning is so easy to measure in the laboratory, one can simply measure innate defensive responses (i.e., SSDRs) to diagnose fear and fear-related memory. This has made fear conditioning one of the major rodent assays of learning, memory and anxiety disorders. Over the last four decades fear conditioning studies have extensively used one of these defensive behaviors, freezing, more than any other response (Anagnostaras et al., 2010; Bouton & Bolles, 1980; Do-Monte et al., 2015; Fanselow & Bolles, 1979; Grewe et al., 2017; Kim & Fanselow, 1992; Kwon et al., 2015; Nader et al., 2000; Roy et al., 2017). Freezing is a common and adaptive defensive behavior as it reduces the likelihood of detection and attack by a predator (Fanselow & Lester, 1988).

However, if rodents have multiple defensive responses, an important theoretical question is what are the conditions that select between different SSDRs (Fanselow, 1997). An influential model of SSDR selection applied to both humans and rodents is Predatory (or Threat) Imminence Continuum theory, which states that qualitatively distinct defensive behaviors are matched to the psychological distance from physical contact with a life-threatening situation (Bouton et al., 2001; Fanselow & Lester, 1988; Mobbs, 2018; Mobbs et al., 2007). Stimuli that model particular points along this continuum elicit behaviors appropriate to that level of predatory imminence. For example, rodents freeze when they detect a predator but show vigorous bursts of activity to contact by the predator (Fanselow & Lester, 1988). The former, labeled post-encounter defense, relates to fear-like states. The latter, referred to as circa-strike defense, relates to panic-like states (Bouton et al., 2001; Perusini & Fanselow, 2015).

According to this account, in fear conditioning experiments the shock US models painful contact with the predator and therefore invariably produces circa-strike activity bursts but not freezing (Fanselow, 1982). On the other hand, stimuli associated with shock such as an auditory CS, models detection of a predator and therefore invariably produces post-encounter freezing as a CR but not activity bursts (Fanselow, 1989).

Recently, two reports challenge this view. Fadok et al. (2017) used a unique two component serial CS consisting of a 10-sec tone followed immediately by a 10-sec white noise ending with a 1 sec shock and found that the initial component (tone) produced freezing, while the second component (noise) produced bursts of locomotion and jumping in mice. Gruene et al. (2015) reported that in rats a tone CS resulted in a similar burst of locomotion, labelled darting. The results were interpreted as a competition between “active” and “passive” defenses. These findings not only challenge the above response selection rule but, also call for a “reinterpretation of rodent fear conditioning studies” because if only one SSDR is measured (e.g., freezing) but the situation is characterized by a different SSDR, fear and fear-related learning may be misdiagnosed (Gruene et al., 2015). Also note that contrary to Predatory Imminence Theory, Gruene et al. (2015) suggested that freezing and darting were competing CRs to the same level of threat (Fanselow, 1989).

Both previous reports concluded that these activity bursts were CRs because they increased over trials during acquisition when CS and US were paired and decreased during extinction when the CS was presented alone (Fadok et al., 2017; Gruene et al., 2015). While these behavioral patterns are certainly properties of a CR, they are not diagnostic of associative learning as these changes could also result from nonassociative processes such as sensitization and habituation (Rescorla, 1967). Additionally, no formal assessment was made of what properties of the CS led to the alternate CRs (e.g., its serial nature, the ordering of the two sounds, or stimulus modality). One subsequent study using this serial conditioning procedure in mice has suggested that this white-noise elicited activity burst is mainly a result of the stimulus salience or intensity of the white noise and does not depend on any particular temporal relation to the US (Hersman et al., 2020). Another recent study using this procedure in rats has suggested that this flight behavior only occurs in context in which fear has been established and is a result of associative processes, but some of the metrics used to score this flight are confounded with any potential freezing prior to the noise presentation (Totty et al., 2021). Therefore, to better understand the associative nature of these flight responses, we embarked on a series of experiments to test these theoretical views and assess the validity of these concerns (Tables 1-4).

**Table 1:** Design of Experiment 1

| **Group** | **Training Treatment: 10 CS-US Pairings**<br>**(5 per day)** | **Testing Treatment**<br>**(5 on one day)** |
| --- | --- | --- |
| 1) Replication | 10-sec Tone→10-sec Noise→1-sec Shock | 10-sec Tone→10-sec Noise |
| 2) CS Duration | 10-sec Noise→shock | 10-sec Noise |
| 3) Stimulus Change | 20-sec Tone→shock | 10-sec Tone→10-sec Noise |

## Results

### Experiment 1

Experiment 1 was conducted as delineated in Table 1 (see Fig. 1 for a schematic representation of the serial conditioned stimulus and the design for training and testing for Experiment 1). We first conducted a nearly exact replication of the conditions used by Fadok et al. (2017), using male and female mice (Replication Group). Briefly, animals received 10 pairings of footshock and the two-component stimulus (10-sec tone followed by 10-sec white noise) over 2 days before being tested on the third day with the two-component stimulus. We scored bursts of locomotion and jumping with a Peak Activity Ratio (PAR; Fanselow et al., 2019) and the number of darts (Gruene et al., 2015). PAR reflects the largest amplitude movement made during the period of interest, while darts reflect the frequency of large movements during the same period (see methods). We included two additional groups in this experiment to test the nature of any observed behaviors. We asked whether any observed behavior occurred to the noise because it was embedded in a serial compound or because of the brevity of the noise (10 sec). For one group, we simply conditioned and extinguished a 10-sec white noise (CS Duration Group). A third group of mice was trained with a 20-sec tone, but tested with the two-component serial compound stimulus (Stimulus Change Group).

**Figure 1.**
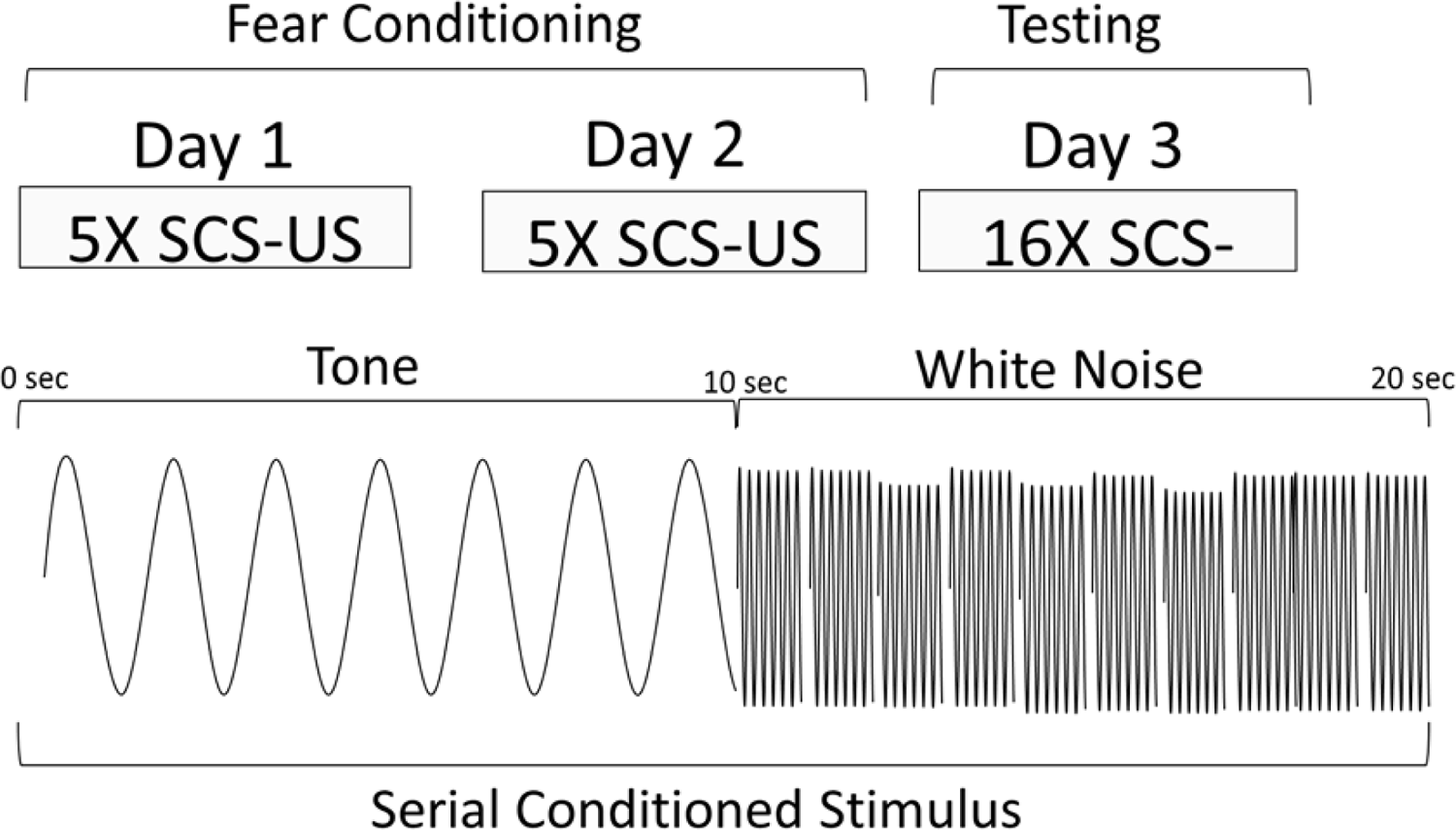
Behavioral design and schematic representation of the serial compound conditioned stimulus (SCS) used for the Replication Group in Experiment 1. During training, animals were given two days each of 5 SCS-US pairings. The SCS consisted of a 10s pure tone (7.5 kHz) followed by a 10s white noise (75 dB). Immediately upon termination of the white noise/SCS, a mild footshock US (1s, 0.9 mA) was delivered. On Day 3, the animals were tested with 16 presentations of the SCS without delivering any shocks.

In a nearly exact replication of the conditions used by Fadok et al. (2017), using male and female mice, we obtained nearly identical results with our Replication Group (Table 1, Fig. S2). For this and all experiments described below, no effects of sex were observed in initial comparisons/ANOVAs (see Discussion). Sex was thus removed as a factor in subsequent statistical analyses. In the replication group, freezing to the initial tone progressively increased over the course of conditioning. At the beginning of training, freezing increased to the white noise but plateaued after a few trials. When freezing plateaued the noise elicited activity bursts, and this pattern maintained throughout acquisition and extinction testing.

Then, we directly asked whether the plateau in freezing and increase in activity that occurred to the noise was because of the brevity of the noise (10-sec) and its close temporal relation to the US. We simply conditioned and extinguished a 10-sec white noise (CS Duration Group) and found that freezing increased linearly during a 10 sec pre-noise period reflecting the acquisition of contextual fear conditioning (Kim & Fanselow, 1992; Fig. S3). During testing the reaction to onset of the white noise was almost a duplicate to what we saw when the noise was embedded in the compound. In other words, activity bursts and darting in no way depended on the use of a serial compound.

To probe the necessity of the compound during acquisition we trained a third group of mice with a 20-sec tone instead of the compound but tested them with the serial compound stimulus (Stimulus Change Group). During these shock-free tests the noise evoked a very similar PAR and darting behavior to when training was with the compound (Fig. 2). What is striking about this finding is that even though the noise was never paired with shock it still evoked an activity burst. These findings strongly implicate nonassociative processes in the activity burst rather than conditioning.

**Figure 2.**
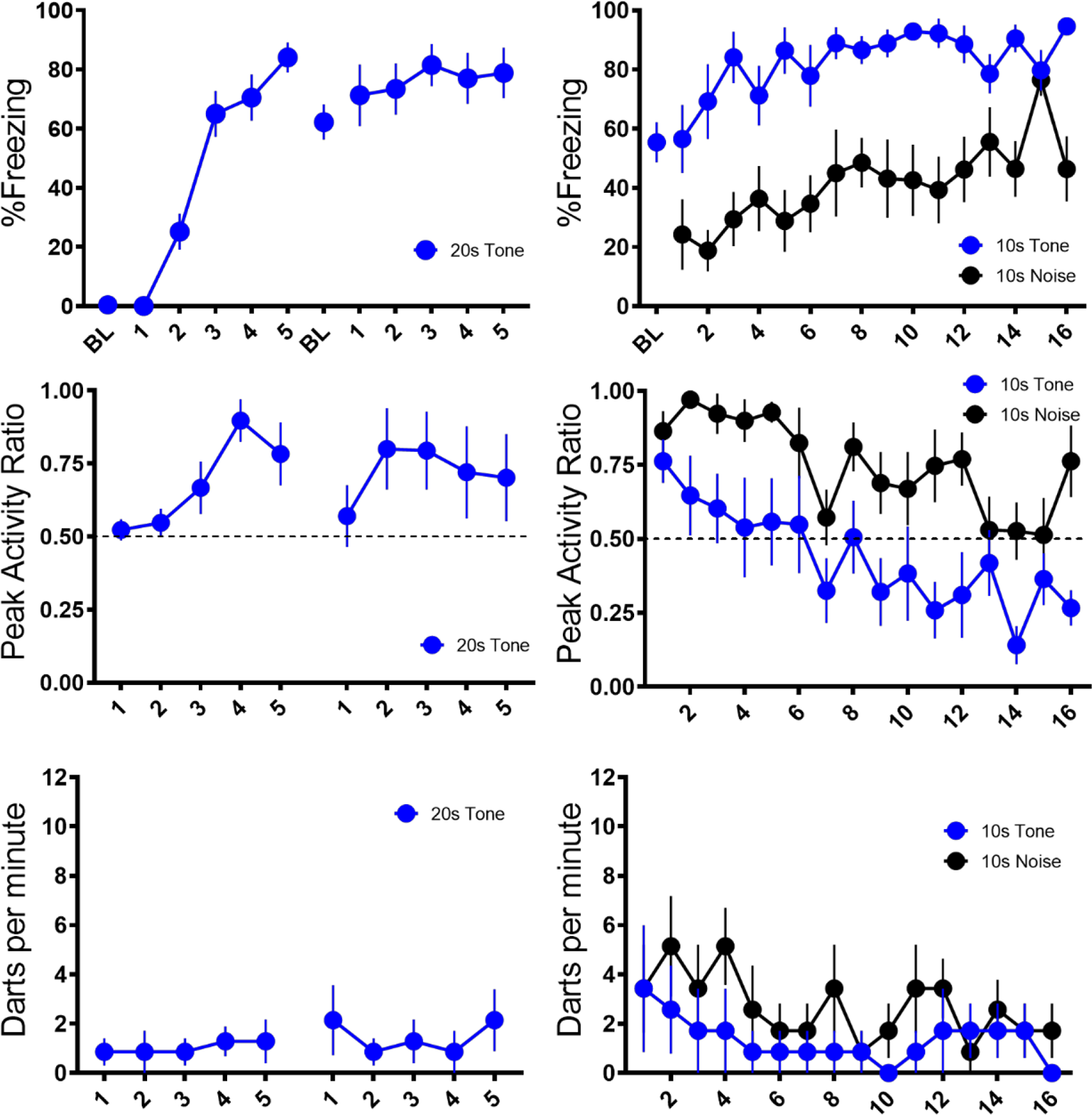
Trial-by-Trial Mean (±SEM) Percent Freezing, Peak Activity Ratio (PAR), and Darts per minute throughout all stimulus presentations during training (left panels) and testing (right panels) for the Stimulus Change Group in Experiment 1.

Overall in Experiment 1, we replicated findings that differential defensive behaviors develop to separate components of a serial CS (Replication Group). This pattern of behavior holds true if the noise is presented by itself during training (CS Duration group), and this pattern of behavior at testing does not require the noise to be present during training (Stimulus Change Group).

Despite differences in behavioral procedures used across acquisition and extinction, we sought to examine any differences in reactivity to the noise during extinction testing between these three groups. We directly analyzed velocity data across the three groups (Fig. 3). We focused on the first four trials of extinction testing as this is when the majority of the darting behavior occurred, and we further narrowed our analyses to the 10s Noise period as all groups received at least the 10s noise at test.

**Figure 3.**
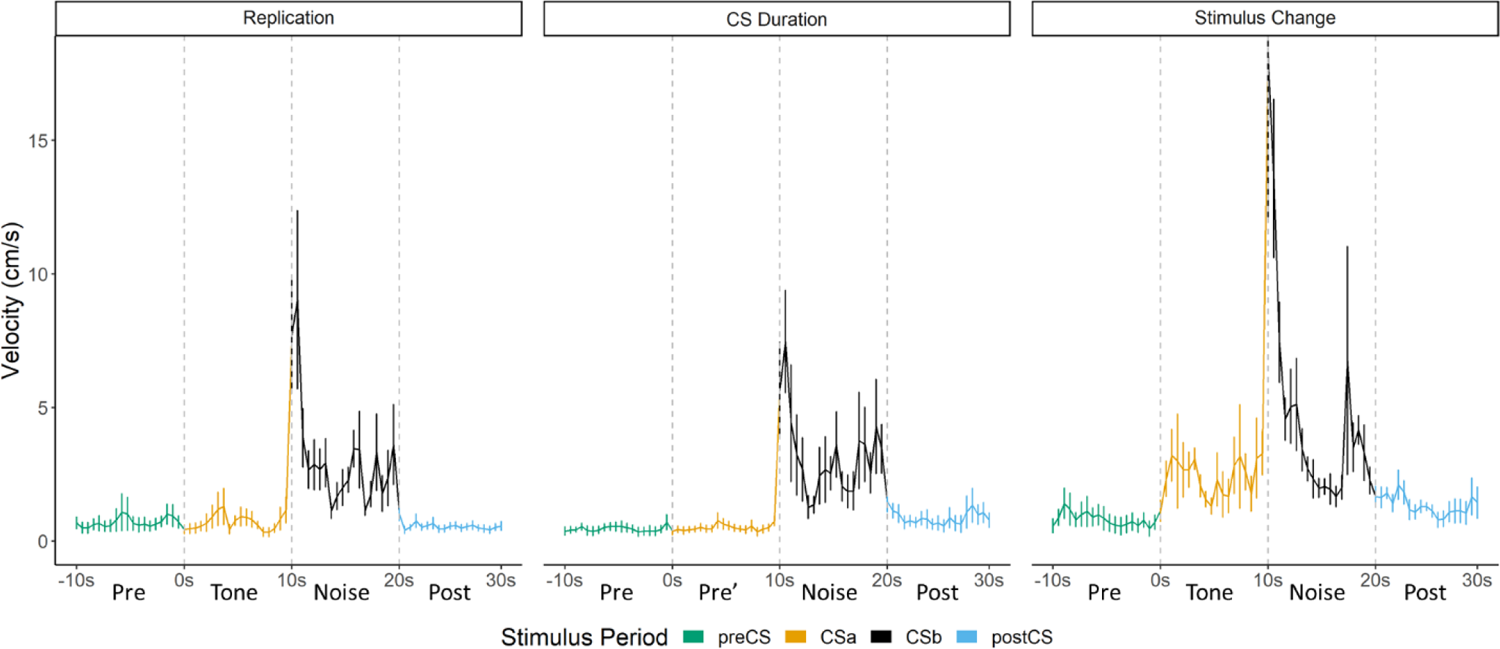
Averaged traces of velocity (cm/s) across the first 4 trials of extinction during testing for Experiment 1. Data is averaged across all animals per group and binned into ∼.5s bins (0.533s) and presented as means plus/minus standard error (Mean ±SE). These within-subject error bars are corrected for between-subject variability using methods as described in Rouder and Morey (2005). During this test, the Replication Group and the Stimulus Change Group received the serial conditioned stimulus (SCS) in which a 10s tone was followed by a 10s noise. The CS Duration group was only tested with a 10s noise.

A mixed model ANOVA revealed a significant effect of Time, [F(19,361)=8.203, p<.001] as well as a Group X Time interaction [F(38,361)=1.497, p=.034]. Generally, velocity peaked during the first bins of the noise period and then quickly decreased to more stable levels. Post-hoc analyses revealed that the Stimulus Change group trended to have the elevated velocity during the first bin of the noise period with trends for higher velocity than the CS Duration group (p=.09) and did have significantly higher velocity than the CS Duration group during the fifth bin (∼2.5 seconds into the noise; p=.04).

While the noise did not need to be within a serial compound stimulus or even need to be presented during training in order to elicit flight, it is worth noting that the strongest noise-elicited flight occurred for the group which received the serial compound stimulus at test and for which the noise was novel at test.

### Experiment 2

The mice that received the 20 sec tone during training but the compound during testing showed darting to the noise embedded in the compound (Figures 2, 3, S3). Since the noise was not paired with the shock, this suggests that the response to the noise was nonassociative.

However, it is possible that during the initial test trials the response to the noise occurred via second-order conditioning as the noise was paired with the previously reinforced tone. This seems unlikely because most darts were seen at the beginning of testing and decreased over the session. A second-order conditioning interpretation suggests the opposite pattern.

Nonetheless, in a second experiment, we included classic controls to directly test for the phenomenon of pseudo-conditioning (Table 2). Pseudo-conditioning is a form of sensitization whereby mere exposure to the US changes behavior to the stimulus used as a CS (Underwood, 1966), and this appears to be what was observed in Experiment 1 (Stimulus Change Group; Fig. 2). Two pseudoconditioned groups of mice simply received the same shock schedule used in the prior study without any auditory stimuli (no CS). A third was merely exposed to the chamber. The final group was a conditioning group that received noise-shock pairings. All groups received tests with the 10 sec noise, except for one of the pseudoconditioning groups that was tested with the tone.

**Table 2:** Design of Experiment 2

| **Group** | **2 Day Training Treatment:** | **Testing Treatment** |
| --- | --- | --- |
| 1) Pseudoconditioned Noise | 10 Shocks<br>(1-mA, 1-sec, 150-210s intertrial interval) | 5 Noise Presentations<br>(10-sec) |
| 2) Pseudoconditioned Tone | 10 Shocks<br>(1-mA, 1-sec, 150-210s intertrial interval) | 5 Tone Presentations<br>(10-sec) |
| 3) No Shock Control | Context exposure Only<br>(17-min & 15-sec per day) | 5 Noise Presentations<br>(10-sec) |
| 4) Noise-Shock Conditioning | 10 Noise (10-sec)→shock pairings | 5 Noise Presentations<br>(10-sec) |

Figures 4 and 5 summarize the test results from Experiment 2 (see Fig. S4 for trial-by-trial data). As would be expected for a CR, freezing to the noise was greatest in the mice that received noise-shock pairings [F(3,28) = 11.76, p<.001]. Significant associative learning was indicated by more noise-elicited freezing in the paired group than the shock-only trained group tested with the noise. Interestingly, the No Shock group that was tested with the noise gradually increased freezing over the course of noise testing (Fig. S4) suggesting that the 75dB noise itself was aversive to the mice and could support some conditioning of freezing (i.e., it was a weak US).

**Figure 4.**
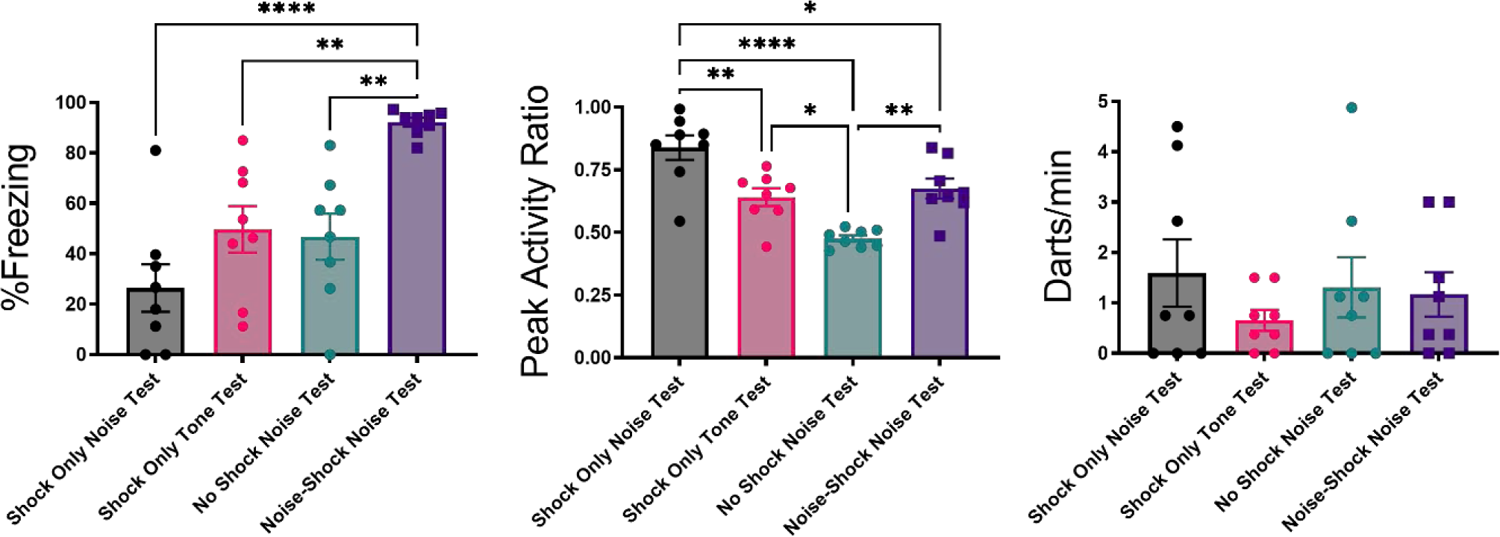
Mean (±SEM) Percent Freezing, Peak Activity Ratio (PAR), and Darting for the test session for Experiment 2. Values are averaged across the 16 trials of extinction during test. *p<.05, **p<.01, ****p<.0001

**Figure 5.**
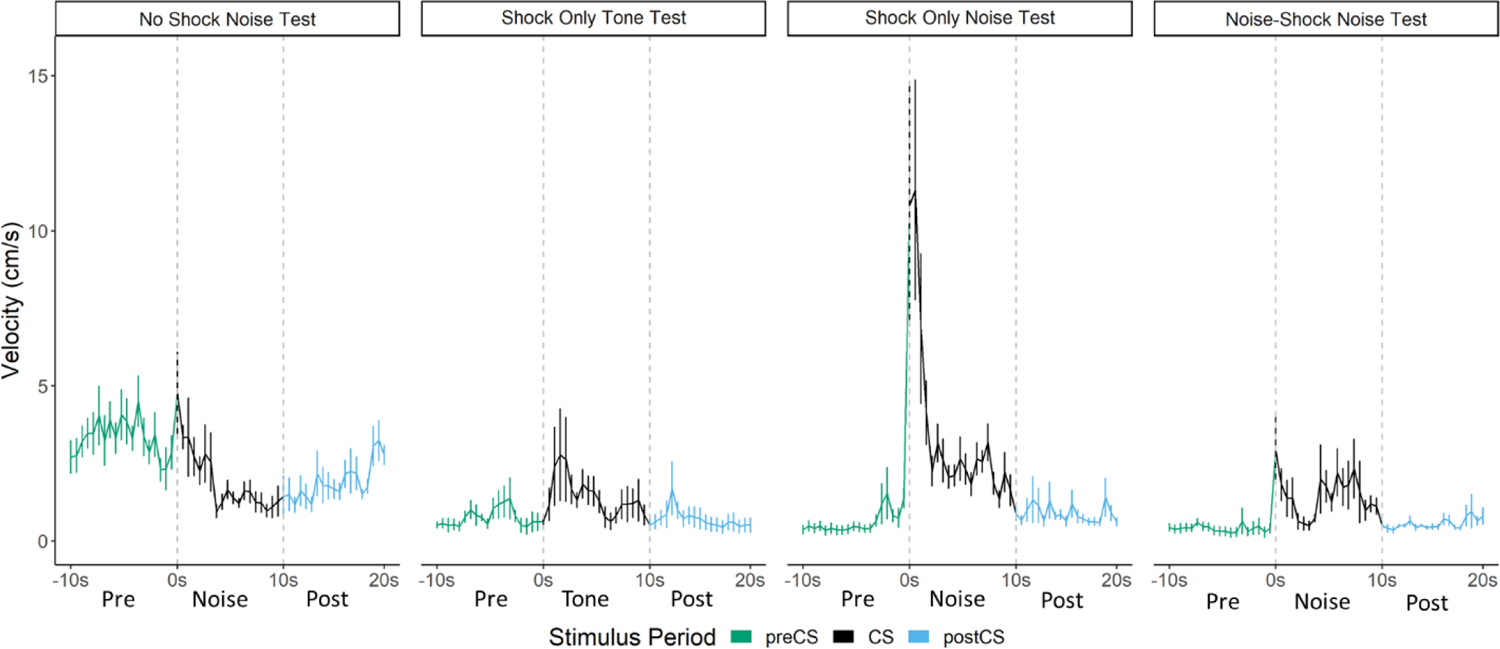
Averaged traces of velocity (cm/s) across the first 4 trials of extinction during testing for Experiment 2. Data is averaged across all animals per group and binned into ∼.5s bins (0.533s) and presented as means plus/minus standard error (Mean ±SE). These within-subject error bars are corrected for between-subject variability using methods as described in Rouder and Morey (2005). During this test, the No Shock-Noise Test, Shock Only-Noise Test, and Noise-Shock Noise Test groups were tested with a 10s noise. The Shock Only-Tone Test group was tested with a 10s tone.

The test session data were very different for activity bursts (Figs. 4 and 5). The greatest PAR occurred in the pseudoconditioned control (shock only during training) that was tested with the novel noise [F(3,28) = 20.085, p<.001]. The pseudoconditioned control tested with the novel noise showed the most darting behavior. Furthermore, these results are supported by a direct analysis of velocity data during the 10s CS period at test (Fig. 5).

A mixed model ANOVA on the averaged velocity measures during the CS period for the first 4 trials of the test session revealed significant effects of Group [F(3,28)=5.796, p=.003] and Time, [F(4.06,113.69)=6.038, p<.001] as well as a Group X Time interaction [F(12.18,113.69)=2.695, p=.003]. Generally, velocity again peaked during the first bins of the noise period and then quickly decreased to more stable levels. Post-hoc analyses revealed that the Shock Only-Noise Test group had the highest velocity during the second bin of the noise period (the first second of the CS) with significantly higher velocity than the No Shock-Noise Test (p=.03), Shock Only-Tone Test (p=.004) and, importantly, the Noise Shock-Noise Test groups (p=.007).

Pseudoconditioning is indicated by more activity during the noise test in the previously shocked mice than the no-shock controls tested with the same noise. Note that for both of these groups the noise was novel during testing so it had no association with shock. Another striking finding is that while the group that received noise-shock training showed an elevated PAR, the level was significantly less than the pseudoconditioning control (p<.001). Not only are activity bursts not conditioned, they are actually suppressed by conditioning! In other words, flight and darting are a result of nonassociative processes and are not CRs.

### Experiment 3

In a third experiment, we included a control group in which the shock and noise were explicitly unpaired to again test for the phenomenon of pseudo-conditioning but in a situation where exposure to the CS is equated during training (Table 3). One group was again a conditioning group that received noise-shock pairings, and one group was again a pseudoconditioned group that only received shocks without any CS. One group received equal numbers of noise and shock presentations but in an explicitly unpaired manner. An additional control group received presentations of only the white noise CS to examine whether or not the CS alone was able to support conditioning and/or activity bursts.

**Table 3:** Design of Experiment 3—Paired vs Unpaired Noise-Shock

| **Group** | **Training Treatment: 10 CS-US Pairings**<br>**(5 per day)** | **Testing**<br>**Treatment**<br>**(2 on one day)** |
| --- | --- | --- |
| 1) Paired Noise-Shock (Conditioning) | 10-sec Noise→1-sec Shock | 10-sec Noise |
| 2) Unpaired Noise/Shock | 10-sec Noise & 1-sec Shock – Unpaired | 10-sec Noise |
| 3) Noise - CS Only | 10-sec Noise | 10-sec Noise |
| 4) Shock Only (Pseudoconditioning) | 1-sec Shock | 10-sec Noise |

Acquisition and test results are summarized in Figures 6 and 7. As seen in the prior experiments, across training freezing to the white noise rises, and then plateaus in the Paired and Unpaired groups, at which point the noise begins to elicit activity bursts. In the CS only group white noise alone supported low, but consistent levels of freezing but in the shocked groups the noise disrupted freezing to the context. During training, the Paired and Unpaired groups showed elevated PAR to the noise [F(3,28)=29.94, p<.001 for Day 1; F(3,28)=75.18, p<.001 for Day 2], and increased darting to the noise [F(3,28)=8.187, p<.001 for Day 1; F(3,28)=22.538, p<.001 for Day 2]. Interestingly, for darting, the Paired group showed elevated responding on Day 2 compared to the Unpaired group (p=.026). During testing, activity bursts (measured as both PAR and darting) to the noise were elevated in all groups which received shock [F(3,28) = 13.35, p<.001 for PAR; F(3,28) = 8.302, p<.001 for darting]. Again, similar to during training, darting was the most elevated in the Paired group on Trial 1 of testing (p=.001).

**Figure 6.**
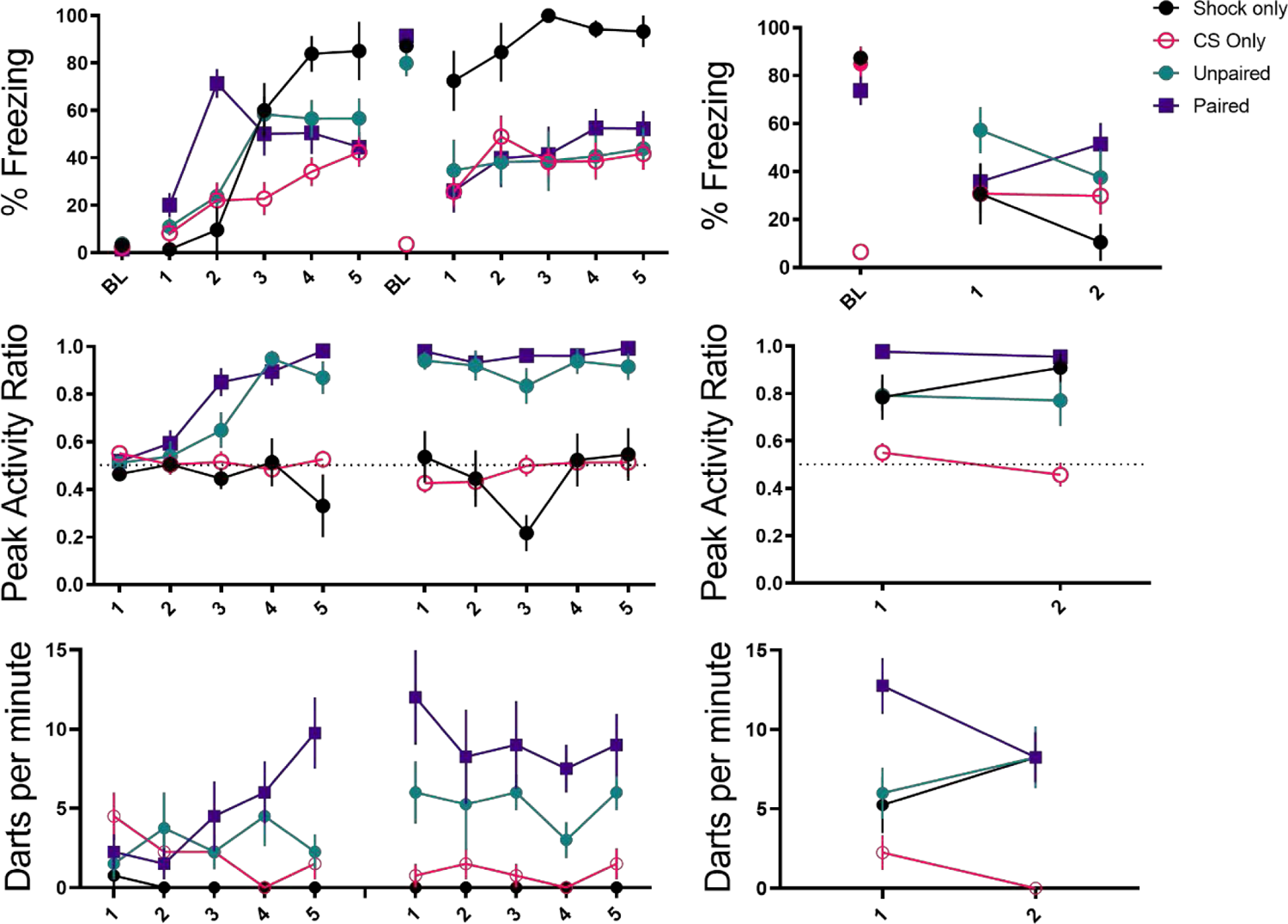
Trial-by-trial Mean (±SEM) Percent Freezing, Peak Activity Ratio (PAR), and Darting per minute throughout all stimulus presentations during training (left panels) and testing (right panels) for Experiment 3.

**Figure 7.**
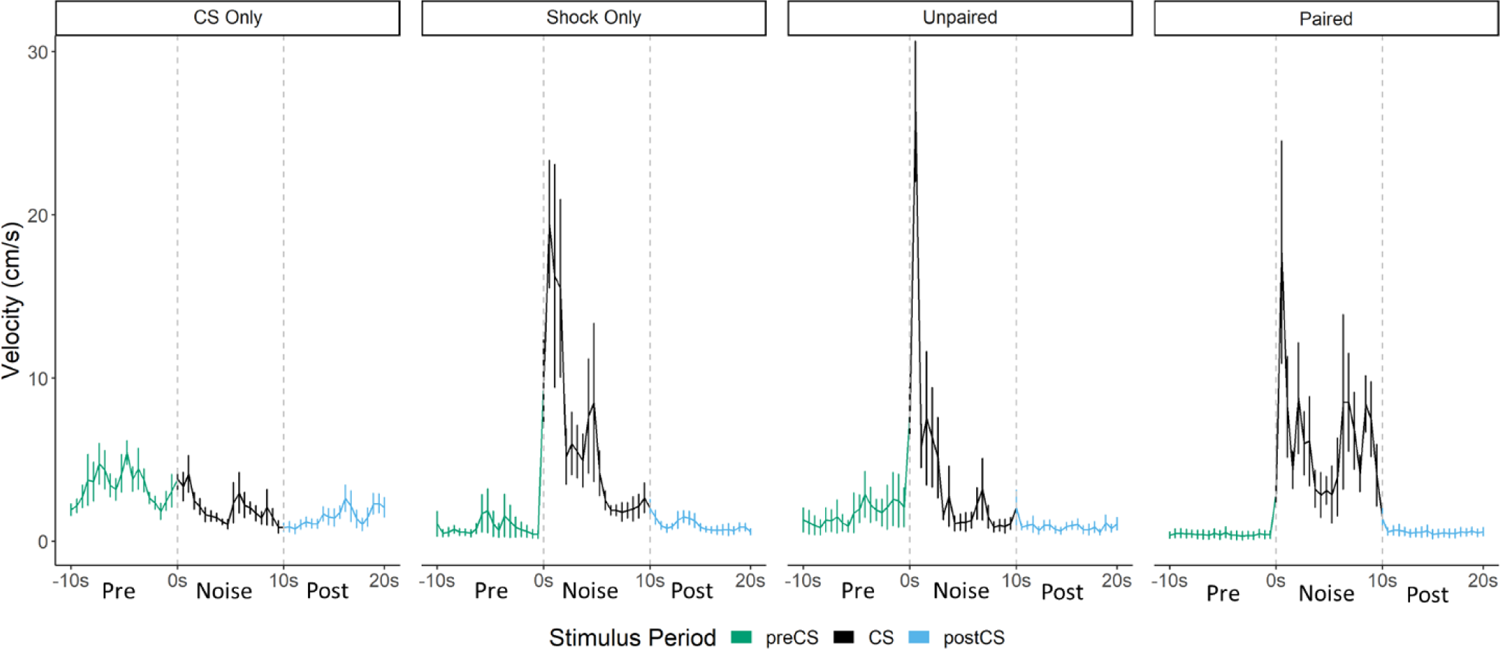
Averaged traces of velocity (cm/s) across 2 trials of extinction during testing for Experiment 3. Data is averaged across all animals per group and binned into ∼.5s bins (0.533s) and presented as means plus/minus standard error (Mean ±SE). These within-subject error bars are corrected for between-subject variability using methods as described in Rouder and Morey (2005). During this test all groups were tested with a 10s Noise CS.

While overall darting was elevated in the Paired group [during acquisition and on the first trial of testing], the velocity traces during testing (Fig. 7) reveal that the magnitude/frequency of the initial activity burst to the noise appears to be reduced in the Paired group, and that increased levels of activity bursts during the latter portion of the CS account for any differences in overall numbers of darts. Indeed, a direct analysis of the velocity data during the 10s Noise CS period at test revealed significant effects of Group [F(3,28)=9.733, p<.001], Time, [F(5.15,144.22)=9.614, p<.001] as well as a Group X Time interaction [F(15.45,144.22)=2.045, p=.02]. Generally, as seen in prior experiments, velocity again peaked during the first bins of the noise period and then quickly decreased to more stable levels. In the Paired group specifically, there is an additional peak of activity in the latter half of the stimulus period. Post-hoc analyses revealed that the Unpaired group had the highest velocity during the first bin of the noise period (the first second) with significantly higher velocity than the CS Only Group (p=.007). Additionally, in the 16^th^ and 17^th^ bins towards the end of the CS period, the Paired Group showed the most activity with significantly higher velocity than the CS Only Group (p=.002 & p=.001), the Shock Only Group (p=.001, p=.02), and the Unpaired Group (p<.001, p=.003)

That pairing noise and shock altered the timing of the activity bursts is an interesting fact worth considering and suggests that pairing noise and shock may have primarily resulted in a conditioned freezing response which in fact competes with/reduces any initial non-associative activity/bursting to the white noise. Taken together, this and the prior experiment using control groups to assess pseudoconditioning reveal that a large portion, if not all, of the noise-elicited activity bursts observed are due to non-associative processes which result in an increase in darting behavior to the noise following shock exposure, regardless of any direct training history of the noise with shock. There does appear to be evidence that pairing noise with shock may further increase or alter the timing of this behavior, but by no means is pairing noise with shock necessary to produce these activity bursts.

### Experiment 4

The experiments thus far have suggested that much of the white-noise-elicited activity bursting is a non-associative process. We have also shown that novelty of the CS at test may increase this noise-elicited activity (Figs. 3 & 4). In a final, fourth experiment, we explicitly tested whether habituation to the white noise stimulus prior to noise-shock training would be able to reduce noise-elicited activity bursts. If increased levels of novelty of the CS are driving noise-elicited activity bursts, then prior habituation should reduce the levels of darting to the noise CS. In this experiment, we had four groups which differed in whether they received an additional two days of habituation to the white noise stimulus (5 noise presentations each day) and whether they received noise-shock pairings during training or just shock only (Table 4). One comparison of particular interest was between the habituated or non-habituated Shock Only groups as these groups would directly compare whether prior experience with the CS would decrease darting at test compared to a group for which the CS was completely novel.

Figure 8 shows the results of Experiment 4 during testing (see Figure S5 for trial-by-trial results for freezing, PAR, and darting across habituation, training, and testing). During the two days of habituation, interestingly, we found that within groups which received habituation, a low level of darting to the white noise alone without any shock decreased across day one [F(4,48) = 2.887, p=.026] and increased by the end of the second day of habituation [F(4,48) = 2.793, p =0.36] (Fig. S5). Concurrently, freezing to the white noise increased over habituation trials, again showing that this white noise stimulus alone can act as a US. It is interesting that darting occurred to the white noise at the start of habituation when the CS was very novel, and at the end of habituation once the white noise alone was able to support some level of fear.

**Table 4:** Design of Experiment 4—Tested the Effect of Habituation to the White Noise

| **Group** | **Habituation**<br>**Treatment: 10 CS**<br>**Exposures (5 per day)** | **Training Treatment: 10**<br>**CS-US Pairings**<br>**(5 per day)** | **Testing Treatment**<br>**(3 on one day)** |
| --- | --- | --- | --- |
| 1) Habituation/Shock Only (H-Shock) | 10-sec Noise | 1-sec Shock | 10-sec Noise |
| 2) Habituation/Paired Noise-Shock (H-Paired) | 10-sec Noise | 10-sec Noise→1-sec Shock | 10-sec Noise |
| 3) Context Exposure/Shock Only (C-Shock) | Context Exposure | 1-sec Shock | 10-sec Noise |
| 4) Context Exposure/Paired Noise-Shock (C-Paired) | Context Exposure | 10-sec Noise→1-sec Shock | 10-sec Noise |

**Figure 8.**
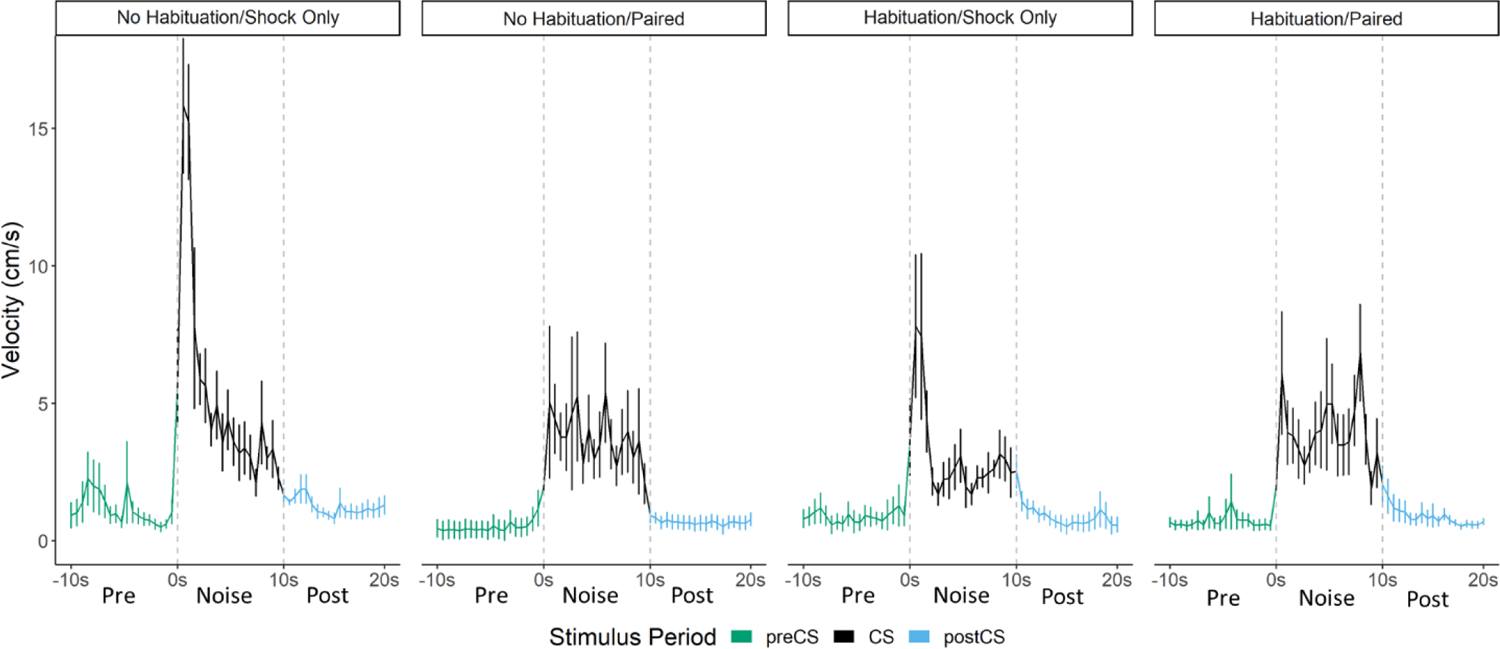
Averaged traces of velocity (cm/s) across 3 trials of extinction during testing for Experiment 4. Data is averaged across all animals per group and binned into ∼.5s bins (0.533s) and presented as means plus/minus standard error (Mean ±SE). These within-subject error bars are corrected for between-subject variability using methods as described in Rouder and Morey (2005). During this test all groups were tested with a 10s Noise CS.

Comparing the two Shock Only groups during test, the noise disrupted freezing more than tone. In this regard noise seems to act like a weak shock US (Fanselow, 1982). Like shock it disrupts freezing (Fig S5) and like shock it supports conditioning of freezing (Fig 6).

Within Paired groups (H-Paired and C-Paired), we found that throughout acquisition and particularly on the second day of training (Fig. S5), prior habituation to the white noise increased freezing [F(1,24)=5.701, p=.025] and decreased noise-elicited darting [F(1,24) = 5.130, p=.033], as predicted if prior exposure to the CS functions to reduce any partially novelty-driven darting. We again saw that freezing to the white noise initially increased during acquisition, but as the darting response begins to become more apparent, freezing decreases to medium levels. At test (Figs. 8 & S5), for freezing, we found a main effect of pairing [F[1,24] = 11.306, p=.003], such that animals who received white noise paired with shock froze more than animals who only received shock during acquisition, again indicative that noise-elicited freezing is a conditional behavior that results from associative learning. For darting behavior, we found a Habituation X Pairing interaction [F(1,28)=4.939, p=.035] such that pairing white noise with shock increased darting within habituated animals (p=.033), and that habituation reduced darting within animals who only received shock during training (p=.045). These results reveal multiple points of interest. First, and as shown in prior experiments, the white noise acts as a US on its own and need not be paired with shock to produce darting at test. Merely experiencing the shock is enough to produce darting to the white noise at test (pseudoconditioning due to sensitization). Furthermore, prior experience with the white noise, through habituation, actually reduced this darting at test. Additionally, in this experiment, we do again show evidence that pairing white noise with shock can further increase darting behavior at test, at least within animals who have already experienced the noise during habituation. Again, as with Experiment 3 (Fig. 7) the timing of the darting response in Paired groups is fundamentally altered compared to Shock Only groups (Fig. 8). The magnitude/frequency of the initial activity burst to the noise appears to be reduced in the Paired groups, and increased levels of activity bursts during the latter portion of the CS account for any differences/increases in overall numbers of darts.

Indeed, a mixed model ANOVA with Pairing, Habituation, and Time as factors on the averaged velocity traces for each trial revealed significant effects of Time [F(56,1568)=17.420, p<.001], a Habituation X Pairing interaction [F(1,28)=4.696, p=.04], and a Pairing X Time interaction [F(56,1568)=3.036, p=.01]. Generally, once again, velocity peaked during the first bins of the noise period and then quickly decreased to more stable levels. As seen in the experiments above, again, this initial peak in velocity was most apparent in the Shock Only groups, with the Paired groups showing an initially smaller peak in velocity. Post-hoc analyses revealed that the Shock Only groups had significantly higher velocity during the first three bins of the noise than the Paired groups (p’s=.02, .03, .005 respectively). Post-hoc analysis on the Pairing X Habituation interaction reveal that within the non-habituated groups, pairing noise and shock significantly reduced the velocity throughout test trials (p<.001). Additionally, within Shock Only groups, habituation reduced the velocity throughout test trials (p<.001). These results are exactly what would be predicted if exposure to the noise CS (through pre-exposure and/or through pairing CS and US) in fact reduces noise-elicited activity bursts and flight/darting behavior, that is, darting is enhanced by novelty.

## Discussion

Prior work reported that contact/pain-related stimuli (e.g., shock) disrupt freezing and provoke panic-like circa-strike defensive behaviors (Fanselow, 1982). The current results suggest a modification of the rules governing a transition between these behavioral states. The rule is that when you are in the post-encounter mode (fear) a sudden change in stimulation, particularly the onset of an intense novel stimulus, can cause an immediate transition to the circa-strike mode (panic). Indeed, the vast majority of the activity bursts/darting behavior occurred at the onset of the stimulus (Figs. 3, 5, 7, 8). The effectiveness of this transition depends on the qualities of the stimulus. Stronger shocks cause a greater disruption of freezing and a longer activity burst, yet the same stronger shocks simultaneously condition more freezing to the prevailing cues (Fanselow, 1982). The current data call for an expansion of this rule to nonnociceptive stimuli. While both tone and noise disrupted ongoing freezing, the noise did so for longer than the tone (Fig. S6) and noise on its own was able to support a minimal level of fear conditioning (Figs. 6, S4, S5). The rule is: when in a state of fear (Post-encounter defense) sudden stimulus change provokes panic-like circa-strike defenses proportional to stimulus intensity and novelty.

As the majority of the experiments presented here and in most prior studies conduct both training and testing in the same context (Fadok et al., 2017; Gruene et al., 2015, Hersman et al., 2020), these animals would already be in a high state of fear or post-encounter defense (from any learned contextual fear during training), thus endowing the presentation of the white noise to be a particularly startling stimulus change which can provoke these panic-like flight responses. Novelty of the stimuli is an important factor and familiarity with the CS during conditioning and/or habituation reduced CS novelty for the test. In the experiments presented here, the mice that received noise-shock pairings and were tested with noise showed lower flight to the noise than mice trained only with shock and then received noise for the first time. Additionally, prior habituation to the noise or experience with the noise during training further reduced noise-elicited flight at test.

Another important factor to consider is the timing of the activity burst with respect to CS and US onset. With poorly timed and sustained conditional responses such as freezing the CR tends to fill the entire CS-US interval and spill over beyond the time of expected US delivery (e.g., Ayres & Vigorito, 1984; Gale et al., 2004). However, shorter duration ballistic responses such as the darting response allow a clearer assessment of when the CR occurs with respect to CS and US delivery and such CRs are expected to anticipate US delivery. Hull (1934) cautioned conditioning researchers that it is important to distinguish true conditional responses from unconditional responses to the CS, which he named alpha responses. These alpha responses occur at the onset of the CS, rather than the time of the expected US. Alpha responses have been most studied with the Pavlovian conditional eyeblink response, where the true CR is well-timed to US delivery (McCormick & Thompson, 1984, Perrett et al,1993). Blinks that occur to CS onset are classified as alpha responses, which are considered to be nonassociative startle responses to the CS and not CRs (e.g., Gerwig et al., 2005; Nation et al., 2017; Schreurs and Alkon, 1990, Woodruff-Pak et al., 1996). Typically, in eyeblink studies alpha responses are excluded from analysis by omitting any responses that occur at the beginning of the CS. Our darting responses almost exclusively occurred at CS onset and there were never any US anticipatory-like responses. Thus, traditional Pavlovian analyses for ballistic CRs would have categorized darting as an unconditional alpha response and not a bona fide CR. Consistent with this analysis is that darting occurred to the noise during the first few trials of the habituation session in Experiment 4 (Fig S5)

Our interpretation that noise unconditionally elicits a ballistic activity burst bears some relationship to the unconditional acoustic startle response. Loud noises will elicit an unconditional startle response that wanes with repeated presentations of that noise (i.e., habituation; e.g., Davis, 1980; Hoffman & Fleshler, 1963; Leaton, 1976). While our 75 dB noise stimulus is less intense than the 98-120 dB noise used in typical acoustic startle studies, we are observing an unconditional noise elicited response that also decreases with habituation (Experiment 4). Furthermore, our data and those of Totty et al. (2021) indicate that these responses require a fearful context in order to occur. Fear is well known to potentiate startle responses (Brown, et al., 1951; Davis, 1989). Perhaps the low intensity noise is below threshold to elicit a startle response on its own, but a fearful context potentiates this response and brings it above threshold. Additionally, there appears to be considerable overlap in the neuroanatomy that supports this circa-strike behavior and fear potentiated startle. Totty et al. (2021) found that inactivation of the Central Nucleus or the Bed Nuclei of the Stria Terminalis disrupts the flight response. These two regions have been shown to be important mediators of fear’s ability to potentiate startle (e.g., Campeau & Davis, 1995; Davis & Walker, 2014). Furthermore, Fadok et al. (2017) reported that it is corticotropin releasing hormone (CRH) expressing cells, but not somatostatin expressing cells, within the Central Nucleus that support flight behavior. Again, there is extensive data implicating CRH and fear potentiated startle (Lee & Davis, 1997).

It is of note that the relationship between startle (circa-strike defense) and freezing (post-encounter defense) was described by Fanselow & Lester (1988) when accounting for how rats rapidly transitioned between these behaviors when a detected predator launches into attack. “It is as if the freezing animal is tensed up and ready to explode into action if the freezing response fails it. This explosive response probably has been studied in the laboratory for over 30 years under the rubric of potentiated startle…It seems that the releasing stimulus for this explosive motor burst is a sudden change in the stimulus context of an already freezing rat (Fanselow & Lester, 1988, p 202).”

Neither Fadok et al. (2017) nor Gruene et al. (2015) included any controls for nonassociative behavior, which is something required in order to conclude that a response is conditional (Rescorla, 1967). Both of these research groups concluded from their single group experiments that flight/darting was a CR because the behavior increased with successive shocks during the shock phase and decreased with shock omission during the test phase, likening these behavioral changes to acquisition and extinction. While acquisition and extinction are characteristics of a CR, learning theorists have never taken these as diagnostic of a CR. For example, increases in responding with successive shocks could arise via sensitization and decreases in responding when shocks are omitted could arise from habituation. Indeed, that is exactly what we believe caused these behavioral changes that we also observed in our study. Shocks, by conditioning fear to the context, sensitize or potentiate the darting response and repeated presentations of the noise alone cause the response to habituate. The behavior of our pseudoconditioning control provides clear evidence of this. Just giving shocks conditioned fear to the context such that when the noise was presented for the first time during test it caused a strong activity burst. The behavior gradually decreased during testing because repeated presentations of the noise led to habituation of this unconditional response.

Given our argument that the flight/darting behavior is nonassociative, Totty et al.’s finding that noise-shock paired rats showed more noise elicited activity burst behavior than rats that had unpaired noise and shock requires additional comment. Since both unpaired and paired rats were exposed to noise during acquisition those exposures could lead to habituation of the unconditional response to the noise. However, it would be expected that habituation would be greater in the unpaired group because pairing a stimulus (noise in this case) with another stimulus (shock in this case) is known to reduce the magnitude of habituation (Pfautz et al., 1978). This reduction in habituation is observed even if the second stimulus is not an unconditional stimulus (Pfautz et al., 1978). Additionally, pairing a habituated stimulus with a US can also cause a return of the habituated alpha response and this loss of habituation is not observed when the two stimuli are not paired (Holland, 1977). Thus, the difference between the paired and unpaired groups reported by Totty et al. (2021) are likely due to differential habituation of the noise during training. This effect of habituation was probably enhanced by Totty et al. including a noise habituation phase prior to training.

Initial reports suggest a sex difference in this noise-elicited flight behavior such that female rats show more of this behavior than males (Gruene et al., 2015). Within each experiment, we found no such sex differences between male and female mice for the PAR and darting measures of flight behavior, and Totty et al. (2021) similarly found no sex differences in such behavior in male and female rats. To further increase the power of such an analysis for sex differences, we pooled all of the groups across the four experiments which received noise-shock pairings. In this analysis, again, we saw no sex differences in flight to the white noise across the two days of acquisition for both PAR [Day1: F(1,29)=.323, p=.58; Day 2: F(1,29)=.507, p=.48], and darting [Day1: F(1,29)=.009, p=.92; Day 2: F(1,29)=3.752, p=.06], and we observed no sex differences across testing to the white noise in extinction for both PAR [F(1,20)=.099, p=.76] and darting [F(1,13)=1.397, p=.258]. Perhaps initial reports of sex differences could be explained by differences in handling and stress provided to females as a result of monitoring estrous phase, a potentially stressful procedure for the animals for which there is not an ideal control in males.

Some have characterized freezing as a passive response that occurs because no other response is available (Blanchard & Blanchard, 1969; Fadok et al., 2017; Gruene et al., 2015; Yu et al., 2016). However, because motion is often the releasing stimulus for predatory attacks it is the best thing for a small mammal like a rat or a mouse to do when a predator is detected and will only be replaced if there is a change consistent with contact (Fanselow & Lester, 1988).

Rodents choose locations in which to freeze such as corners or objects (thigmotaxis) (Grossen & Kelley, 1972). The current data show that the freezing rodent also prepares to react to sudden stimulus change. There is nothing passive about it.

## Methods and Materials

### Subjects

Subjects for all experiments included 120 C57BL/6NHsd mice (Experiment 1, n=24; Experiment 2, n=32; Experiment 3, n=32; Experiment 4, n=32), aged 9-11 weeks of age and purchased from Envigo. This C57BL/6NHsd strain was chosen to match that of Fadok et al. (2017). Each group consisted of 4 male and 4 female mice. A necessary/powerful group sample size of 8 was calculated based both on years of data in our lab which suggests n=8 is sufficient to detect such behavioral differences in fear conditioning studies and on the recent articles in the literature which are using this procedure. Mice were group-housed four per cage on a 12-hr light/dark cycle with ad libitum access to food and water. Across each experiment, mice in each cage were randomly assigned to one of the groups, ensuring that every group had a representative from each cage to avoid any cage effects. All experiments were conducted during the lights-on phase of the cycle. Animals were handled for 5 days prior to the start of experiments. Subjects were all treated in accordance with an approved protocol from the Institutional Animal Care and Use Committee at the University of California-Los Angeles following guidelines established by the National Institute of Health.

### Apparatus and Stimuli

All experiments were conducted in standard MedAssociates fear conditioning chambers (VFC-008; 30.5 x 24.2 x 21 cm), controlled by Med Associates VideoFreeze software (Med Associates, St. Albans, VT). For each experiment, the same context was used for training and testing (see Discussion). The context was wiped down between each mouse with 70% isopropanol and 3 sprays of 50% Windex were added to the pans below the shock grid floors to provide an olfactory cue/context. The US consisted of a 1 second 0.9 mA scrambled shock delivered through a MedAssociates shock scrambler (ENV-414S). Each of the CSs were delivered using a MedAssociates speaker (ENV-224AM-2). The tone was 7.5kHz. Both the tone and the white noise were 75dB inside the chamber. The inter-trial interval varied between 150 seconds and 210 seconds with an average length of 180 seconds.

### Design and Procedure

Mice were handled for 5 days for approximately 1 minute per day prior to beginning the experiment. At the beginning of each day of the experiment, mice were transported in their home cages on a cart to a room adjacent to the testing room and allowed to acclimate for at least 30 minutes. Mice were individually placed in clean empty cages on a utility cart for transport from this room to the testing room and promptly returned to their home cages after the session was over. These transport cages were wiped down with StrikeBac in between trials/sessions.

Experiment 1 was conducted as delineated in Table 1 (see Fig. 1 for a schematic representation of the serial conditioned stimulus and the design for training and testing for Experiment 1). The Replication group was trained on each of the two days with 5 presentations of a 10 second tone immediately followed by a 10 second noise, which was immediately followed by a 1 second shock. On Day 3 it was then tested with 16 presentations of a 10 second tone immediately followed by a 10 second noise. These parameters were chosen to match those of Fadok et al (2017) except that we did not include a session of unreinforced CS preexposure prior to conditioning as such treatment is known to reduce conditioned behavior (Lubow & Moore, 1959; we did add such a treatment to Experiment 4 as an experimental factor). The CS Duration group was trained on each of the two days with 5 presentations of a 10 second noise, which was immediately followed by a 1 second shock. It was tested with 16 presentations of the 10 second noise. The Stimulus Change group was trained on each of the two days with 5 presentations of a 20 second tone immediately followed by a 1 second shock. It was tested with 16 presentations of a 10 second tone immediately followed by a 10 second noise (i.e., the compound used in the replication group). Two mice were excluded from this study due to experimenter error, one female in the Replication group and one female in the Stimulus Change group.

Experiment 2 was conducted as delineated in Table 2. The Pseudoconditioned Noise and Pseudoconditioned Tone groups were trained on each of the two days with 5 presentations of a 1-sec shock without any sound using the same schedule for shocks as Experiment 1. The No Shock Control was merely allowed to explore the context for the same length of time as the other groups without receiving any shock or auditory stimuli throughout the two days of acquisition. The final Noise-Shock Conditioning group was trained on each of the two days with 5 presentations of a 10-sec noise, which was immediately followed by a 1-sec shock. As Experiment 1 revealed that similar behavior was observed in groups which received compound stimulus-shock pairings or just noise-shock pairings, we utilized simple noise-shock pairings in this and some of the following experiments to more specifically assess the associative nature of any white noise-driven behavior. All groups received tests with 16 presentations of the 10-sec noise in extinction, except for one of the pseudoconditioning groups that was tested with the 10-sec tone.

Experiment 3 was conducted as delineated in Table 3. The Paired Noise-Shock (Conditioning) group was trained on each of the two days with 5 presentations of a 10 second noise, which was immediately followed by a 1 second shock. The Unpaired Noise/Shock group was presented with the same number and length of noise and shocks, but they were explicitly unpaired in time. The Noise-CS Only group received 5 presentations of a 10 second noise without receiving any shocks on each of the two days. The Shock Only (Pseudoconditioning) group received 5 presentations of a 1 second shock on each of the two days. As the main behavioral responses and differences between groups occurred primarily in the first few trials of the previous experiments, and in order to more readily complete all of the testing within one day’s light cycle, for this and the following experiments we reduced the number of test trials presented to the animals. Thus, at test for this experiment, all groups received two presentations of a 10 second noise.

Experiment 4 was conducted as delineated in Table 4. Prior to training with shock, all groups underwent 2 days of additional training with either habituation to the white noise or merely exposure to the context. The habituated groups, Habituation/Shock Only (H-Shock) and Habituation/Noise-Shock Pairing (H-Paired), were trained on each of the two days with 5 presentations of a 10-second noise, while the two non-habituated groups, Context Exposure/Shock Only (C-Shock) and Context Exposure/Noise-Shock Pairing (C-Paired) received only equivalent exposure to the context. The following two days, and as in the Experiments above, all groups received 10 footshocks. The Paired groups (H-Paired and C-Paired) were trained on each of the two days with 5 presentations of a 10-second noise, followed immediately by a 1-second footshock. The Shock Only groups (H-Shock and C-Shock) were trained on each of the two days with only 5 presentations of a 1-second footshock. At test, all groups received 3 presentations of the 10-second noise.

### Data and Statistics and Analysis

Freezing behavior for Experiments 1-3 was scored using the near-infrared VideoFreeze scoring system. Freezing is a complete lack of movement, except for respiration (Fanselow, 1980). VideoFreeze allows for the recording of real-time video at 30 frames per second. With this program, adjacent frames are compared to provide the grayscale change for each pixel, and the amount of pixel change across each frame is measured to produce an activity score. We have set a threshold level of activity for freezing based on careful matching to hand-scoring from trained observers (Anagnostaras et al., 2010). The animal is scored as freezing if they fall below this threshold for at least a 1-sec bout of freezing.

For Experiment 4, due to a technical error, videos for the first 4 days of the experiment could not be accurately assessed for freezing behavior using VideoFreeze. Therefore, we alternatively measured and scored freezing behavior using EthoVision. Briefly, videos were converted to MPEG, as described above, and analyzed using the Activity Analysis feature of Ethovision. Thresholds for freezing were again determined to match hand-scoring from trained observers.

Two different measures of flight were used. We scored bursts of locomotion and jumping with a Peak Activity Ratio (PAR); Fanselow et al., 2019) and the number of darts (Gruene et al., 2015). To determine PAR, we took the greatest between frame activity score during a period of interest (e.g., the first 10 s of CS presentation = During) and calculated a ratio of that level of activity to a similar score derived from a preceding control period of equal duration (e.g., 10 s before presentation of the tone = PreStim) of the form During/(During + PreStim). With this measure, a 0.5 indicates that during the time of interest there was no instance of activity greater than that observed during the control period (PreStim). PARs approaching 1.0 indicate an instance of behavior that far exceeded baseline responding. This measure reflects the maximum movement the animal made during the period of interest.

Darting was assessed as in Gruene et al. (2015). Video files from VideoFreeze were extracted in Windows Media Video format (.wmv) and then converted to MPEG-2 files using Any Video Converter (AnvSoft, 2018). These converted files were then analyzed to determine animal velocity across the session using EthoVision software (Noldus), using a center-point tracking with a velocity sampling rate of 3.75 Hz. This velocity data was exported, organized, and imported to R (R Core Team, 2018). Using a custom R code (available as source code 1), darts were detected in the trace with a minimum velocity of 22.9 cm/s and a minimum interpeak interval of 0.8 s. The 22.9 cm/s threshold was determined by finding the 99.5th percentile of all baseline velocity data analyzed, prior to any stimuli or shock, and this threshold was validated to match with manual scoring of darts, such that all movements at that rate or higher were consistently scored as darts. See Figure S1 for representative traces of velocity across Day 1 of acquisition for a mouse in the Replication group of Experiment 1. The PAR measure reflects the maximum amplitude of movement, while the dart measure reflects the frequency of individual rapid movements.

Trial-by-trial Measures of freezing and flight were analyzed with a repeated measure multifactorial analysis of variance (ANOVA) and post hoc Tukey tests. Baseline freezing, and overall responding which were collapsed across session when appropriate, were analyzed with a univariate ANOVA test. To directly compare each groups’ activity and the magnitude of any flight behaviors during extinction testing, velocity data was binned into .533s bins and subsequently analyzed using repeated measures ANOVA in R. Whenever violations of sphericity were found, the Greenhouse-Geisser correction was used to produce corrected degrees of freedom and p-values. Significant effects and interactions were followed up with simple main effects and Bonferroni-corrected pairwise t-tests. A value of p<.05 was the threshold used to determine statistical reliability. For all experiments described above, no effects of sex were observed in initial comparisons/ANOVAs. Sex was thus removed as a factor in subsequent statistical analyses.

---

**Figure S1.**
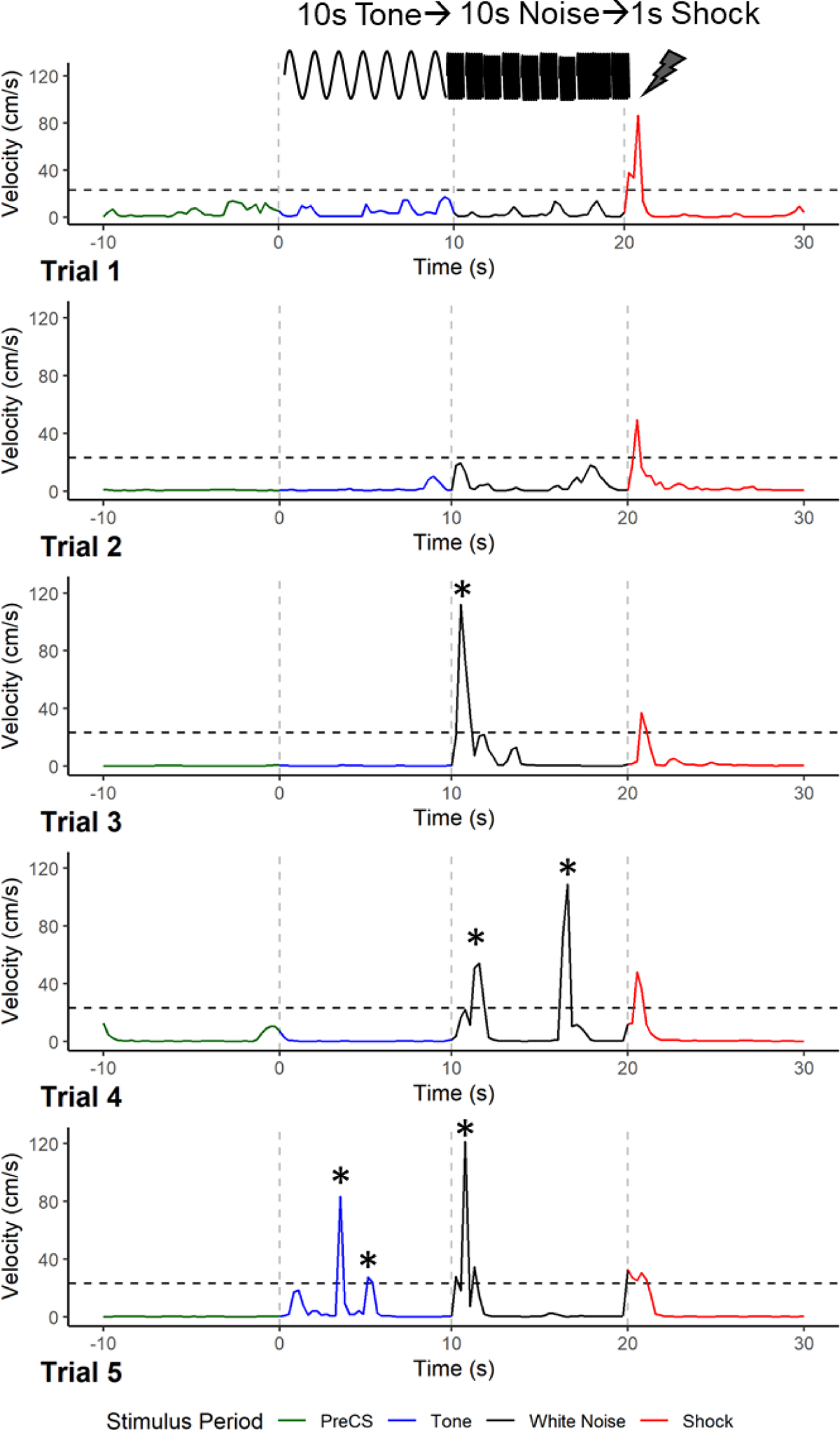
Example traces of velocity (cm/s) measurements obtained via EthoVision across five trials on the first day of training for a mouse in the Replication Group of Experiment 1. Vertical dotted lines denote stimulus onset times and the horizontal dotted line is the threshold for scoring behavior as a dart (22.9 cm/s). Darting episodes are marked with an *.

**Figure S2.**
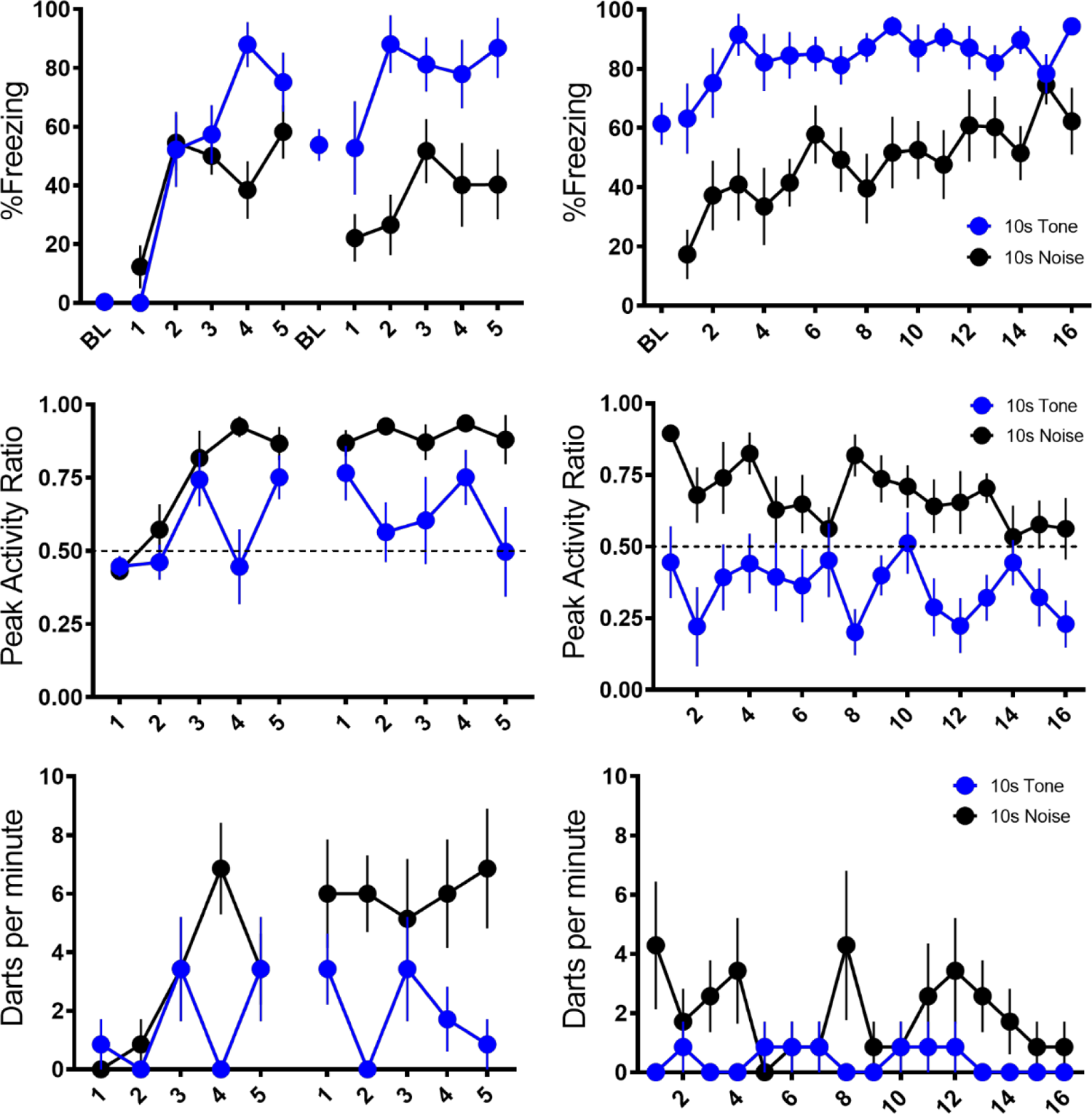
Mean (±SEM) Percent Freezing, Peak Activity Ratio (PAR), and Darts per minute throughout training (left panels) and testing (right panels) for the Replication Group of Experiment 1. Responding during the tone is represented with filled in grey circles, responding during the noise is represented with filled in black circles.

**Figure S3.**
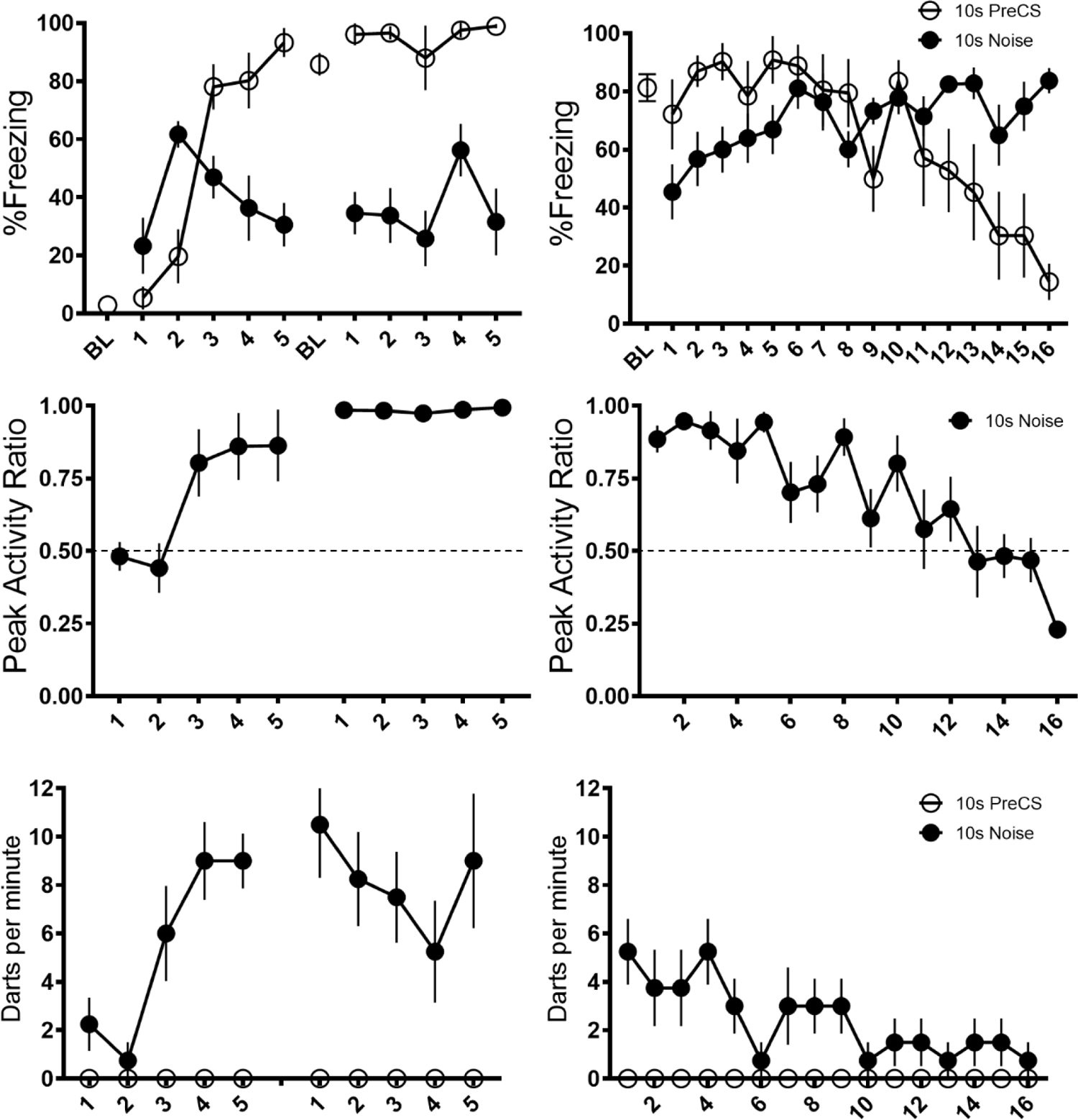
Mean (±SEM) Percent Freezing, Peak Activity Ratio (PAR), and Darts per minute throughout training (left panels) and testing (right panels) for the CS Duration Group of Experiment 1. Responding during the 10s preCS period is represented with open circles, responding during the noise is represented with filled in black circles.

**Figure S4.**
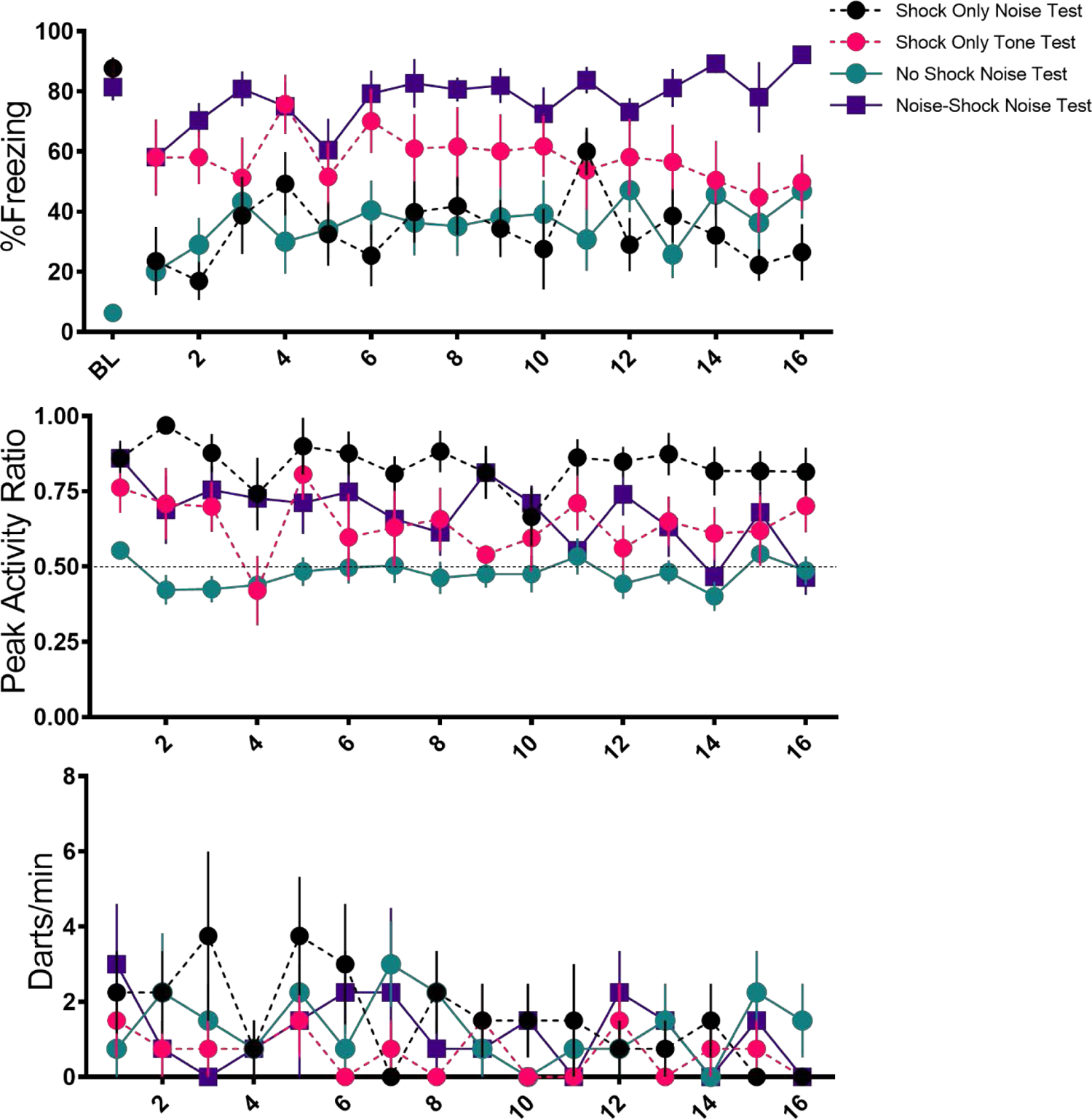
Trial-by-Trial Mean (±SEM) Percent Freezing, Peak Activity Ratio (PAR), and Darts per minute throughout 16 trials of testing for Experiment 2.

**Figure S5.**
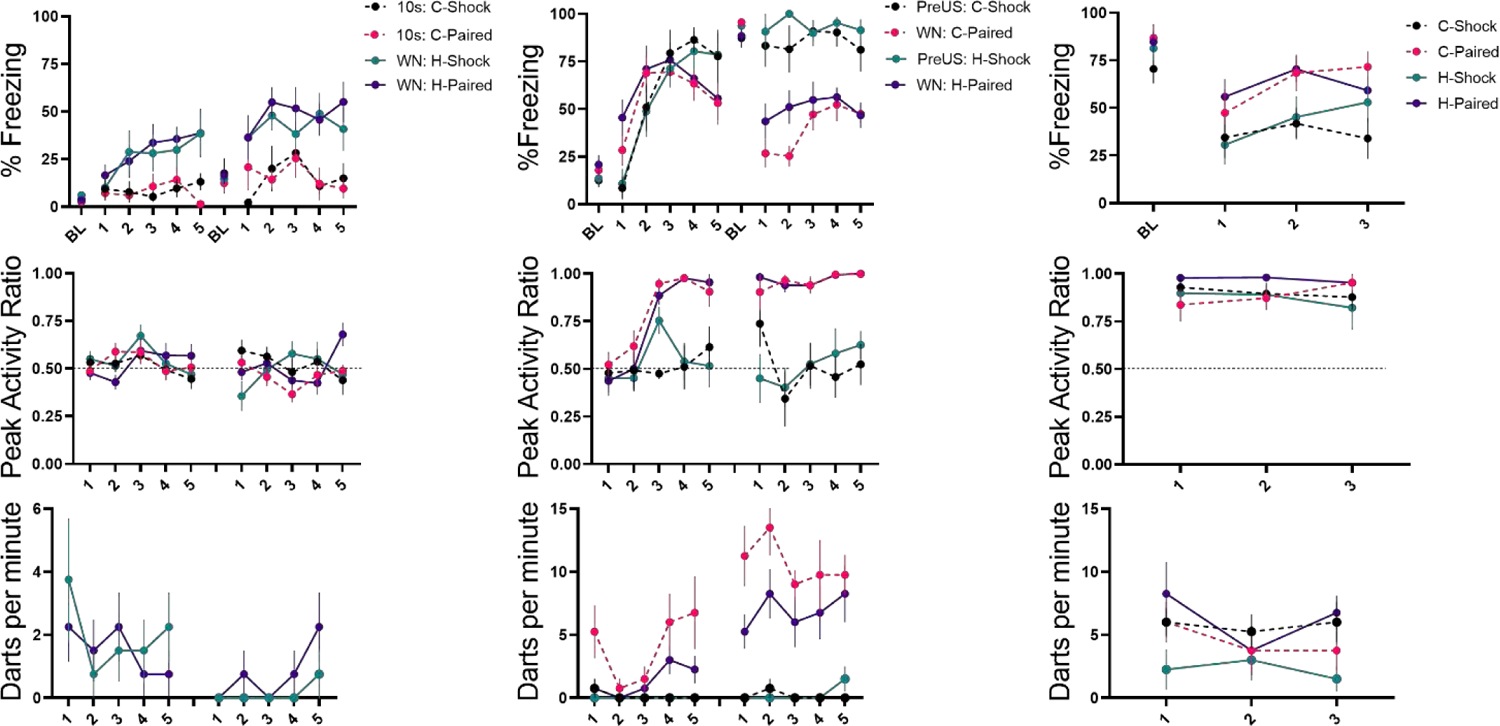
Trial-by-trial Mean (±SEM) Percent Freezing, Peak Ratio (PAR), and Darting per minute throughout all stimulus presentations during habituation (left panels), training (middle panels), and testing (right panels) for Experiment 4.

**Figure S6.**
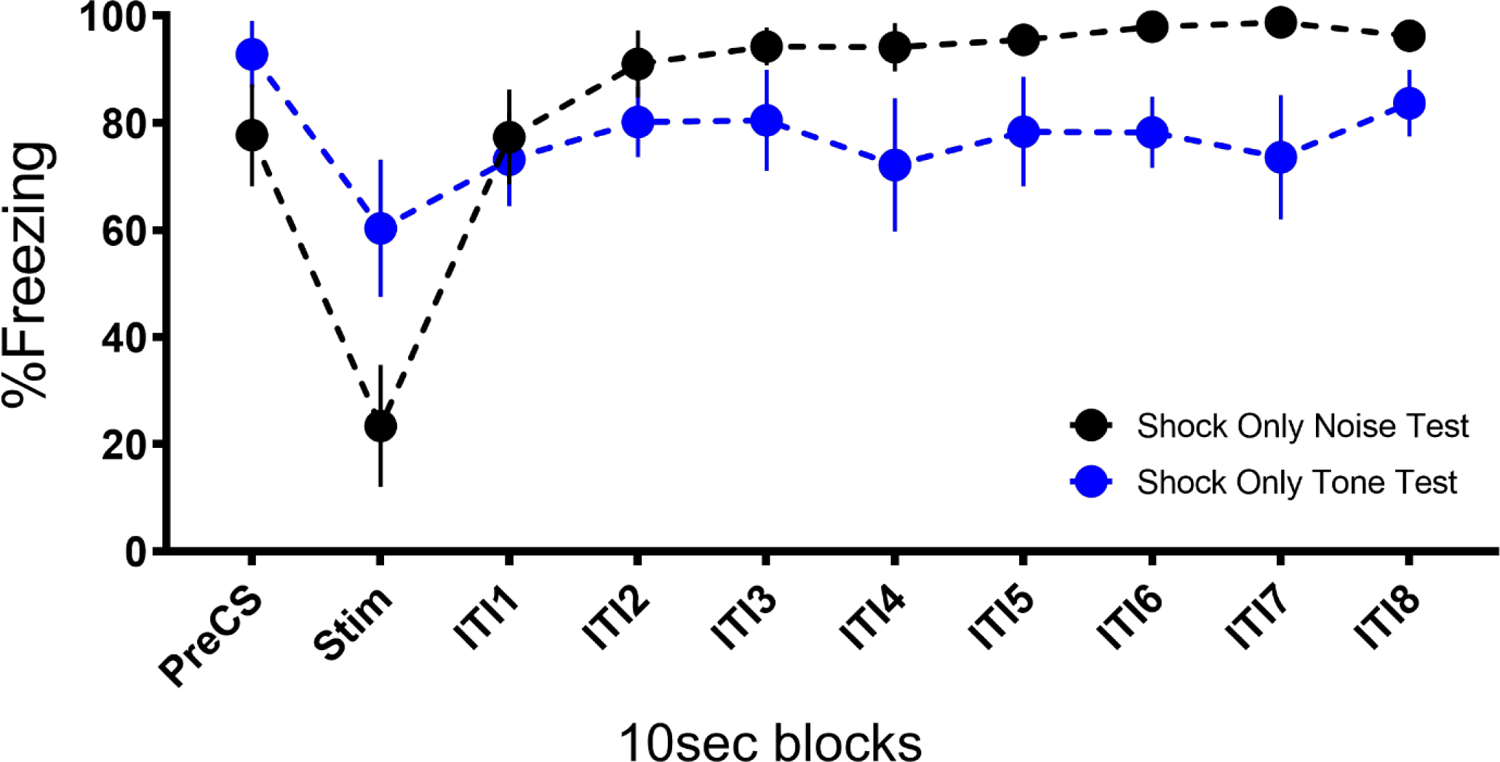
Mean (±SEM) Percent Freezing during extinction/testing for Experiment 2 showing that the occurrence of the stimuli at test disrupt freezing to the context and that the noise disrupts freezing to a greater extent than the tone.

